# Co-cultivation of *Saccharomyces cerevisiae* strains combines advantages of different metabolic engineering strategies for improved ethanol yield

**DOI:** 10.1101/2023.07.04.547682

**Authors:** Aafke C.A. van Aalst, Igor S. van der Meulen, Mickel L.A. Jansen, Robert Mans, Jack T. Pronk

**Author notes:** Corresponding author: Jack Pronk.

## Abstract

Glycerol is the major organic byproduct of industrial ethanol production with the yeast *Saccharomyces cerevisiae*. Improved ethanol yields have been achieved with engineered *S. cerevisiae* strains in which heterologous pathways replace glycerol formation as the predominant mechanism for anaerobic re-oxidation of surplus NADH generated in biosynthetic reactions. Functional expression of heterologous phosphoribulokinase (PRK) and ribulose-1,5-bisphosphate carboxylase (RuBisCO) genes enables yeast cells to couple a net oxidation of NADH to the conversion of glucose to ethanol. In another strategy, NADH-dependent reduction of exogenous acetate to ethanol is enabled by introduction of a heterologous acetylating acetaldehyde dehydrogenase (A-ALD). This study explores potential advantages of co-cultivating engineered PRK-RuBisCO-based and A-ALD-based strains in anaerobic bioreactor batch cultures. Co-cultivation of these strains, which in monocultures showed reduced glycerol yields and improved ethanol yields, strongly reduced the formation of acetaldehyde and acetate, two byproducts that were formed in anaerobic monocultures of a PRK-RuBisCO-based strain. In addition, co-cultures on medium with low acetate-to-glucose ratios that mimicked those in industrial feedstocks completely removed acetate from the medium. Kinetics of co-cultivation processes and glycerol production could be optimized by tuning the relative inoculum sizes of the two strains. Co-cultivation of a PRK-RuBisCO strain with a *Δgpd1 Δgpd2* A-ALD strain, which was unable to grow in the absence of acetate and evolved for faster anaerobic growth in acetate-supplemented batch cultures, further reduced glycerol formation but led to extended fermentation times. These results demonstrate the potential of using defined consortia of engineered *S. cerevisiae* strains for high-yield, minimal-waste ethanol production.

## 1. Introduction

*Saccharomyces cerevisiae* is extensively used for industrial production of ethanol, the largest-volume product of industrial biotechnology (1). Yeast-based ethanol production predominantly occurs in the United States of America and Brazil, using hydrolyzed corn starch or cane sugar, respectively, as main feedstocks (1). These ‘first-generation’ industrial ethanol production processes can reach ethanol yields on sugar of up to 92% of the theoretical maximum (2). The remaining sugar is mainly converted to biomass and glycerol (3, 4). Anaerobic fermentation of glucose or sucrose starts by oxidation of these sugars to pyruvate via the Embden-Meyerhof glycolysis, yielding ATP and NADH (5, 6). Reduction of pyruvate by pyruvate decarboxylase and NAD^+^-dependent alcohol dehydrogenase re-oxidizes the NADH formed in glycolysis and completes the conversion of sugars into ethanol and carbon dioxide (7).

In *S. cerevisiae*, formation of biomass and glycerol are coupled via redox-cofactor balances. Biomass formation leads to a net production of NADH. In anaerobic cultures, this ‘surplus’ NADH can neither be re-oxidized by mitochondrial respiration nor by the redox-cofactor-balanced pathway for ethanol production. Instead, anaerobic yeast cultures rely on the NADH-dependent reduction of the glycolytic intermediate dihydroxyacetone-phosphate to glycerol-3-phosphate (8, 9). This reaction is catalyzed by Gpd1 and Gpd2 (10, 11) and followed by the hydrolysis of glycerol-3-phosphate to glycerol and phosphate by Gpp1 and Gpp2 (12). As approximately 4% of the potential ethanol yield in industrial processes is estimated to be lost to glycerol (4), multiple metabolic engineering strategies have focused on reducing glycerol formation by engineering of yeast redox metabolism (13).

Introduction into *S. cerevisiae* of phosphoribulokinase (PRK) and ribulose-1,5-bisphosphate carboxylase/oxygenase (RuBisCO), the two key enzymes of the Calvin cycle, enables a non-oxidative bypass of glycolysis (14, 15). Ribulose-5-phospate can be generated via the non-oxidative pentose pathway, thereby creating a redox-cofactor neutral bypass from sugars to 3-phosphoglycerate. This concept has been implemented and optimized to allow for fast-growing low-glycerol-producing *S. cerevisiae* strains with an up to 10% higher ethanol yield on glucose in fast-growing anaerobic laboratory cultures (0.29 h^-1^; (15)).

In industrial ethanol fermentation, the specific growth rate decreases as the ethanol concentration reaches inhibitory levels and/or non-sugar nutrients are depleted (16). A recent study showed that, in slow-growing anaerobic chemostat cultures (0.05 h^-1^), a PRK-RuBisCO strain produced up to 80-fold more acetaldehyde and 30-fold more acetate than a reference strain (17). This production of acetaldehyde and acetate was attributed to an *in vivo* overcapacity of the key enzymes of the PRK-RuBisCO bypass. Reduction of the copy number of the expression cassette for RuBisCO led to lower acetaldehyde and acetate production in slow-growing cultures and a corresponding increase in ethanol yield (17). The production of acetaldehyde and acetate was further decreased by reducing PRK activity by lowering protein abundance by a C-terminal extension of PRK, or by expressing the spinach *prk* gene from the growth-rate-dependent *ANB1* promoter (17, 18). However, the resulting strains still showed trade-offs in performance at low and high specific growth rates, which illustrated the challenges involved in tuning the activity of engineered pathways under dynamic conditions.

An alternative redox-engineering strategy is based on the expression of a heterologous gene encoding acetylating acetaldehyde dehydrogenase (A-ALD). Together with native acetyl-CoA synthetase and alcohol dehydrogenase, A-ALD can catalyse the NADH-dependent reduction of exogenous acetate to ethanol (19). In the presence of acetate, anaerobic cultures of engineered A-ALD-expressing strains carrying a deletion in *GPD2*, which encodes one of the two *S. cerevisiae* isoenzymes of glycerol-3-phosphate dehydrogenase, show reduced glycerol yields and improved ethanol yields on glucose (20). When also *GPD1* is deleted, A-ALD strains can even grow anaerobically in the presence of acetate without producing glycerol (19, 21). However, feedstocks for first-generation ethanol processes typically contain insufficient acetate to reach the same ethanol yield improvement as PRK-RuBisCO strains (22).

Although only few industrial biotechnology processes are currently based on defined co-cultures, there is a growing interest in the potential advantages of this approach (23–25). Co-cultivation of PRK-RuBisCO and A-ALD-based strains may, in theory, prevent accumulation of acetaldehyde and acetate generated by PRK-RuBisCO strains while limiting reduced fermentation rates and increased glycerol yields of A-ALD-based strains when acetate in growth media is depleted (Fig. 1). To evaluate this strategy, the present study explores co-cultivation of PRK-RuBisCO and A-ALD-based strains in anaerobic batch cultures on glucose, grown in the presence and absence of acetate. In addition to an A-ALD-containing strain in which only *GPD2* is deleted and which can therefore still grow in the absence of exogenous acetaldehyde or acetate, co-cultures with an A-ALD strain lacking both glycerol-3-phosphate dehydrogenase isoenzymes are investigated.

**Fig 1.**
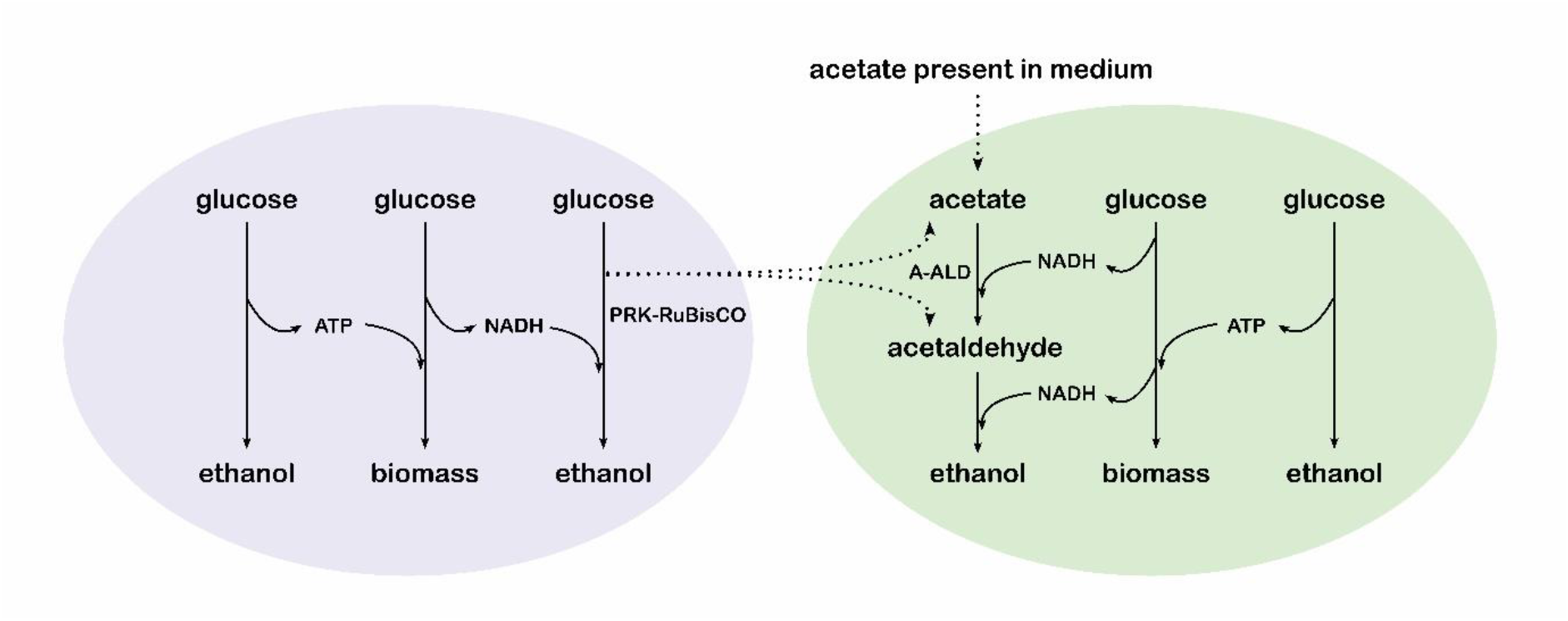
Schematic representation of NADH balances in anaerobic co-cultures of a PRK-RuBisCO-containing S. cerevisiae strain (left) and an A-ALD-containing strain (right). In both strains, re-oxidation of NADH generated during anaerobic biomass formation is coupled to ethanol production. Dotted arrows indicate use of the by-products acetaldehyde and acetate, generated by the PRK-RuBisCO-dependent strain, as electron acceptors for NADH reoxidation by the A-ALD-dependent strain.

## 2. Results

### 2.1 Acetaldehyde and acetate as byproducts of PRK-RuBisCO-containing S. cerevisiae strains

In previous studies, growth and product formation by PRK-RuBisCO-containing *S. cerevisiae* strains were studied in anaerobic bioreactor batch cultures on 20 g L^-1^ glucose (15, 17). In such cultures, fast exponential growth continues until glucose is almost completely consumed. To capture some of the dynamics in large-scale processes, anaerobic bioreactor batch cultures of the reference strain IME324 and the PRK-RuBisCO-based strain IMX2736 (*Δgpd2* non-ox PPP↑ *prk* 2× *cbbm groES groEL*; non-ox PPP↑ indicates integration of the overexpression cassettes for the non-oxidative pentose phosphate pathway genes *RPE1*, *TKL1*, *TAL1*, *NQM1*, *RKI1* and *TKL2*, (17)) were grown on synthetic medium with 50 g L^-1^ glucose. In these cultures, both strains showed an initial exponential growth phase with a specific growth rate of 0.3 h^-1^, after which the growth rate gradually declined to approximately 0.15 h^-1^ (Fig. 2, Fig. S1). These growth dynamics mimicked those in industrial fermentation processes for ethanol production from corn starch hydrolysates (16). Due to these changing growth rates, stoichiometries of glucose consumption and product formation could not be assumed constant throughout batch cultivation. Overall stoichiometries were therefore calculated from measurements on the first and final two time points of cultivation experiments.

Consistent with previous studies on PRK-RuBisCO-based *S. cerevisiae* strains, the glycerol yield on glucose of strain IMX2736, grown anaerobically on 50 g L^-1^ glucose, was 73% lower than that of the reference strain, while its ethanol yield on glucose was 7% higher (p = 0.037, Table 1). In these cultures of strain IMX2736, yields of the byproducts acetate and acetaldehyde on glucose were 0.030 and 0.018 mol (mol glucose)^-1^, respectively (Table 1). This acetaldehyde yield was two-fold higher than reported for anaerobic cultures of strain IMX2736 on 20 g L^-1^ glucose, in which the specific growth rate remained 0.3 h^-1^ throughout batch cultivation (17). Since acetaldehyde production by anaerobic chemostat cultures of PRK-RuBisCO-based strains increases at low specific growth rates (17), the higher acetaldehyde yield of batch cultures grown on 50 g L^-1^ glucose was attributed to their declining specific growth rate.

In earlier batch experiments with PRK-RuBisCO-based *S. cerevisiae* strains grown on 20 g L^-1^ glucose, biomass yields on glucose were approximately 5% higher than in cultures of reference strains (15, 17). Those observations were consistent with lower ATP costs for NADH regeneration via the RuBisCO bypass than via the native glycerol pathway (13). In cultures grown on 50 g L^-1^ glucose, biomass yields of strains IME324 and IMX2736 were not significantly different. Absence of a higher biomass yield of strain IMX2736 in these cultures may be related to the accumulation of 4.1 ± 0.4 mmol L^-1^ acetaldehyde and 7.8 ± 0.1 mmol L^-1^ acetate in the culture broth. Combined, formation of these metabolites already accounted for a loss of ca. 6 mmol L^-1^ glucose, while loss of acetaldehyde via the gas phase (26), toxicity effects of acetaldehyde and weak-acid uncoupling by acetate may affect biomass yield even further (27–29).

**Table 1.** Key physiological parameters of anaerobic bioreactor batch cultures of *S. cerevisiae* strains IME324 (reference), IMX2736 *(Δgpd2* non-ox PPP↑ *prk* 2× *cbbm groES*/*groEL*), IMX2503 (*Δgpd2 Δald6 eutE*) and IMS1247 *(Δgpd1 Δgpd2 Δald6 eutE*, evolved). Non-ox PPP↑ indicates integration of the overexpression cassettes for *RPE1, TKL1, TAL1, NQM1, RKI1 and TKL2*. Cultures were grown on synthetic medium with 50 g L^-1^ of glucose, with or without addition of 5 mM acetate, at pH 5 and at 30 °C and sparged with a 90:10 mixture of N_2_ and CO_2_. Y indicates yield, subscript x denotes biomass. Acetate and acetaldehyde concentrations indicate values in the culture broth, measured at the end of the cultivation experiments. Yields were calculated using the average of the first two and last two sampling points. Degree-of-reduction balances (61) were used to verify data consistency. Values represent averages ± mean deviations of measurements on independent duplicate cultures for each combination of strain and medium. n.d., not determined; n.a., not applicable (acetate consumption instead of acetate production).

| Strain name | IME324<sup>a</sup> |  | IMX2736 |  | IMX2503<sup>b</sup> | IMS1247 |
| --- | --- | --- | --- | --- | --- | --- |
| Relevant genotype | reference | | $\Delta gpd2 prk 2x cbbm$ | | $\Delta gpd2 \Delta ald6 eutE$ | $\Delta gpd1 \Delta gpd2 \Delta ald6 eutE$ |
| Initial acetate concentration | 0 mM | 5 mM | 0 mM | 5 mM | 5 mM | 5 mM |
| Y<sub>biomass/glucose</sub> (g<sub>x</sub> g<sup>-1</sup>) | 0.084 ± 0.000 | 0.084 ± 0.000 | 0.082 ± 0.005 | 0.081 ± 0.001 | 0.085 ± 0.000 | 0.100 ± 0.001 |
| Y<sub>ethanol/glucose</sub> (mol mol<sup>-1</sup>) | 1.49 ± 0.02 | 1.55 ± 0.01 | 1.60 ± 0.00 | 1.64 ± 0.01 | 1.58 ± 0.00 | 1.78 ± 0.02 |
| Y<sub>acetaldehyde/glucose</sub> (mol mol<sup>-1</sup>) | <0.001 | n.d. | 0.018 ± 0.001 | n.d. | n.d. | n.d. |
| Y<sub>glycerol/glucose</sub> (mol mol<sup>-1</sup>) | 0.169 ± 0.006 | 0.150 ± 0.000 | 0.046 ± 0.002 | 0.041 ± 0.001 | 0.125 ± 0.000 | <0.001 |
| Y<sub>acetate/glucose</sub> (mol mol<sup>-1</sup>) | 0.012 ± 0.002 | n.a. | 0.030 ± 0.002 | 0.017 ± 0.002 | n.a. | n.a. |
| mmol glycerol per g<sub>x</sub> | 11.2 ± 0.3 | 9.9 ± 0.0 | 3.1 ± 0.0 | 2.8 ± 0.1 | 8.2 ± 0.0 | 0.1 ± 0.0 |
| Final concentration acetaldehyde (mM) | 0.1 ± 0.0 | n.d. | 4.1 ± 0.4 | n.d. | n.d. | n.d. |
| Final concentration acetate (mM) | 2.8 ± 0.3 | 4.4 ± 0.1 | 7.8 ± 0.1 | 9.3 ± 0.5 | <0.01 | <0.01 |
| Electron recoveries | 99-101 | 99-99 | 98-98 | 99-100 | 99-99 | 101-102 |

<sup>a</sup>data on strain IME324 and IMX2503 were taken from (22).

**Fig. 2.**
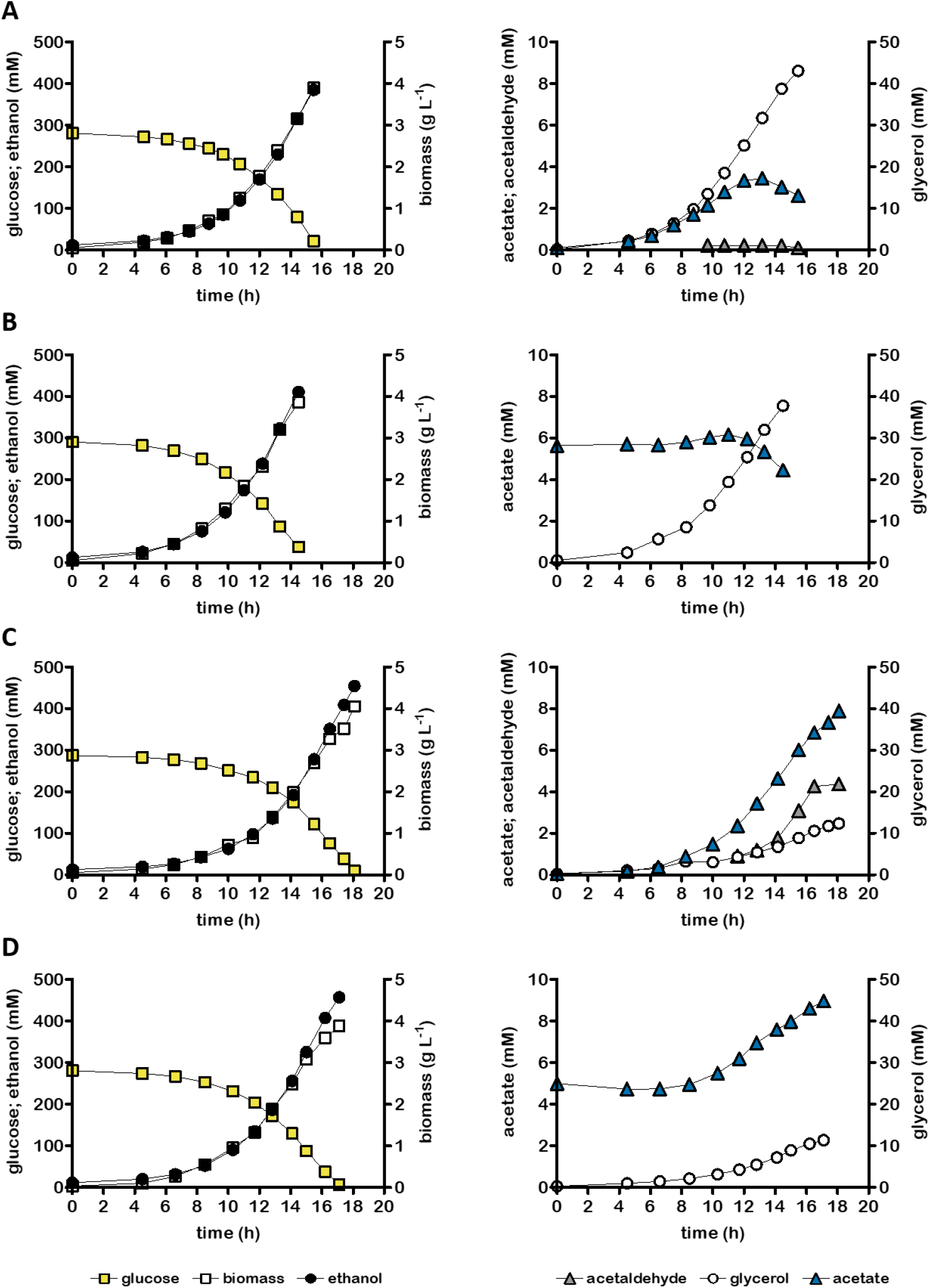
Growth, glucose consumption and product formation of anaerobic bioreactor batch cultures of individual *S. cerevisiae* strains, grown on SM with 50 g L^-1^ glucose (panels A and C) or on SM with 50 g L^-1^ glucose and 5 mmol L^-1^ acetate (panels B and D). Panels show data for *S. cerevisiae* strains IME324 (reference strain, A and B) and IMX2736 *(Δgpd2* non-ox PPP↑ *prk* 2× *cbbm groES/groEL*, C and D). Non-ox PPP↑ indicates integration of the overexpression cassettes for *RPE1*, *TKL1*, *TAL1*, *NQM1*, *RKI1* and *TKL2*. Representative cultures of independent duplicate experiments are shown, corresponding replicate of each culture is shown in Fig. S2.

### 2.2 Anaerobic co-cultivation of PRK-RuBisCO-based and A-ALD-based S. cerevisiae strains on acetate-containing medium

First-generation feedstocks for ethanol production such as corn mash can contain up to 20 mmol L^-1^ acetate (30–32) while glucose concentrations can reach 300 g L^-1^ (33, 34). To mimic these glucose-to-acetate ratios, anaerobic bioreactor batch cultures were grown on 50 g L^-1^ glucose and 5 mmol L^-1^ acetate (Table 1, Fig. 2). Under these conditions, the reference strain IME324 consumed approximately 1.6 mmol L^-1^ acetate (Table 1), probably reflecting its conversion to the biosynthetic precursor molecule acetyl-Coenzyme A (35). In contrast, the PRK-RuBisCO-containing strain IMX2736 produced 2.8 mmol L^-1^ acetate. In these acetate-supplemented anaerobic cultures, the PRK-RuBisCO strain showed a 2.4% higher ethanol yield than in cultures grown without acetate supplementation (p = 0.011, Table 1), which may reflect an increased ATP demand caused by mild weak-acid uncoupling by acetate (27).

Introduction of a heterologous acetylating acetaldehyde dehydrogenase (A-ALD) enables anaerobic *S. cerevisiae* cultures to re-oxidize ‘surplus’ NADH from biosynthetic reactions by NADH-dependent reduction of exogenous acetate to ethanol (19, 22). In anaerobic batch cultures on 50 g L^-1^ glucose and 5 mmol L^-1^ acetate, the A-ALD-containing strain IMX2503 (*Δgpd2 Δald6 eutE*; (22)), had already converted all acetate after 12 h, when 67% of the glucose was still available (Fig. 4A). Moreover, the rate of acetate consumption already declined before this time point (Fig. 4A). Consequently, over the whole process, glycerol formation via Gpd1 was the predominant mechanism for reoxidizing ‘surplus’ NADH in strain IMX2503 and its glycerol yield on glucose of was only 17% lower than that of the reference strain IME324 (p < 0.001, Table 1).

The PRK-RuBisCO strain IMX2736 (*Δgpd2* non-ox PPP↑ *prk* 2× *cbbm groES/groEL;* (17)) does not depend on acetate for glycerol-independent NADH cofactor balancing and its byproducts acetate and acetaldehyde could potentially be used as electron acceptors for the A-ALD-strain IMX2503 (Fig. 1). We therefore investigated whether co-cultivation of these two strains could combine low-glycerol fermentation with complete conversion of acetate. Two unique SNPs in strain IMX2736, on Chromosome 11 (location 331347) and Chromosome 15 (location 912014), allowed for estimation of the cell ratio of strains IMX2503 and IMX2736 in co-cultures by counting reads containing and lacking these SNPs in whole-genome sequence data (Fig. 3).

When strains IMX2503 (A-ALD) and IMX2736 (PRK-RuBisCO) were inoculated at a ratio of 1.4:1 (IMX2503:IMX2736) in medium containing 50 g L^-1^ glucose and 5 mmol L^-1^ acetate, acetate was completely consumed when, after 14 h, ca. 50% of the glucose was still unused (Fig. 4B). When the initial abundance of the A-ALD-based strain was decreased by changing the inoculum ratio to 0.8:1 (IMX2503:IMX2736), complete consumption of glucose and acetate almost coincided (Fig. 4C). As a consequence, acetate limitation of A-ALD strain was delayed and the glycerol yield of the co-culture on glucose was 20% lower than at the higher inoculum ratio (p < 0.001, Table 2). Glycerol yields in co-cultures grown at inoculum ratios of 0.8 and 1.4 were 49% and 34% lower than in corresponding monocultures of the reference strain IME324 (Tables 1 and 2). Ethanol yields of the consortium cultures with an inoculum ratio of 0.8 were not significantly different from that of a monoculture of the PRK-RuBisCO strain IMX2736 and 6% higher than that of monocultures of the reference strain IME324 (p = 0.015, Tables 1 and 2). Biomass yields on glucose of these co-cultures were 6.7% higher than those of a monoculture of the PRK-RuBisCO strain IMX2736 grown on the same medium (p = 0.002, Tables 1 and 2). The higher biomass yield of the consortium cultures is likely to reflect consumption of acetaldehyde and acetate by the A-ALD strain.

**Fig. 3.**
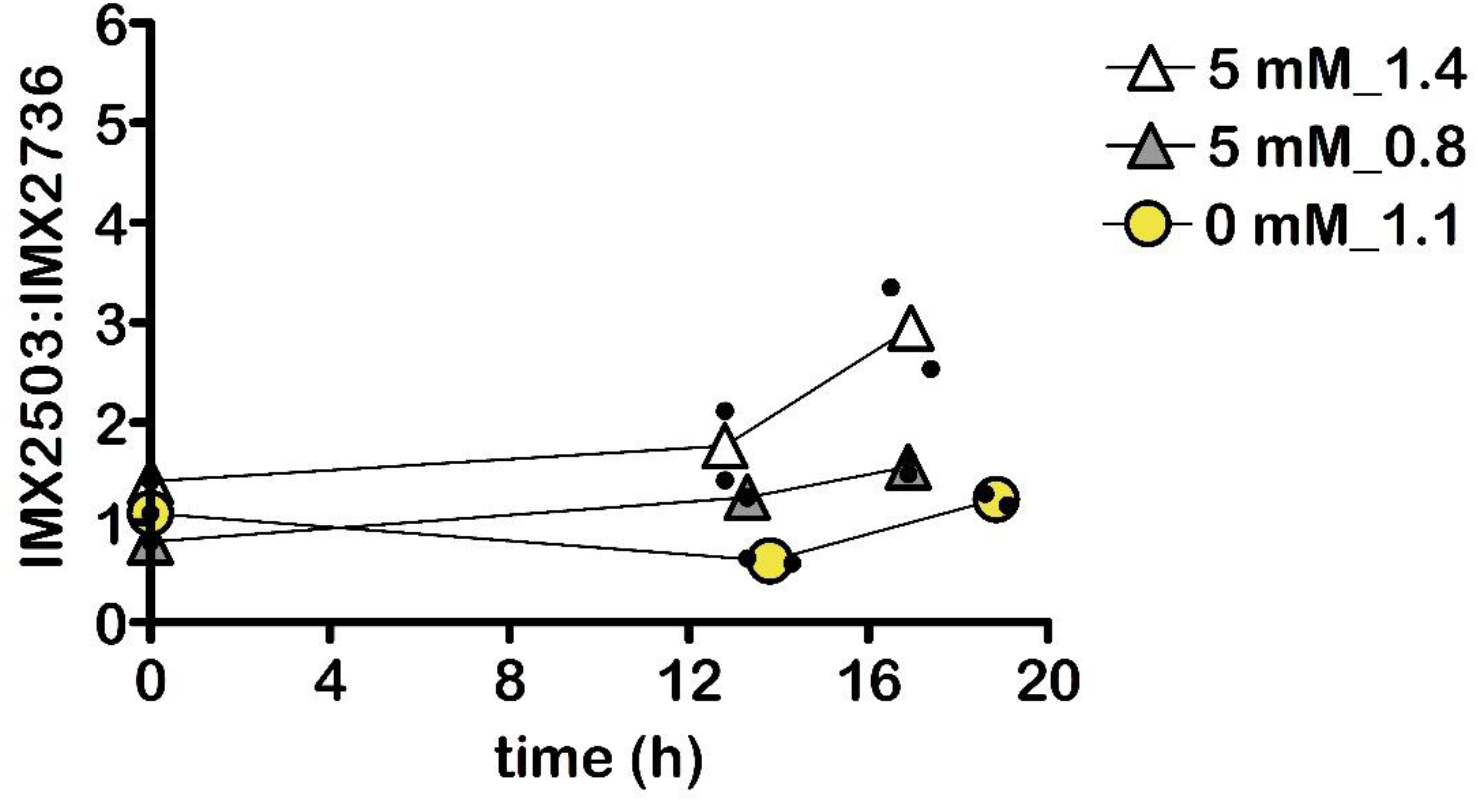
Ratio of *S. cerevisiae* strains IMX2503 *(Δgpd2 Δald6 e*utE) relative to IMX2736 *(Δgpd2*, non-ox PPP↑, *prk*, 2× *cbbm*, *groES, groEL*) in anaerobic bioreactor batch co-cultures on synthetic medium containing 50 g L^-1^ glucose, with or without the addition of 5 mM acetate. Ratio was calculated based on whole genome sequencing. Non-ox PPP↑ indicates the integration of the overexpression cassettes for *RPE1*, *TKL1*, *TAL1*, *NQM1*, *RKI1* and *TKL2*. Values represent means and individual values of measurements on independent batch duplicate cultures. Cultures on 50 g L^-1^ glucose and 5 mM acetate were inoculated at two different ratios of the two strains.

**Fig. 4.**
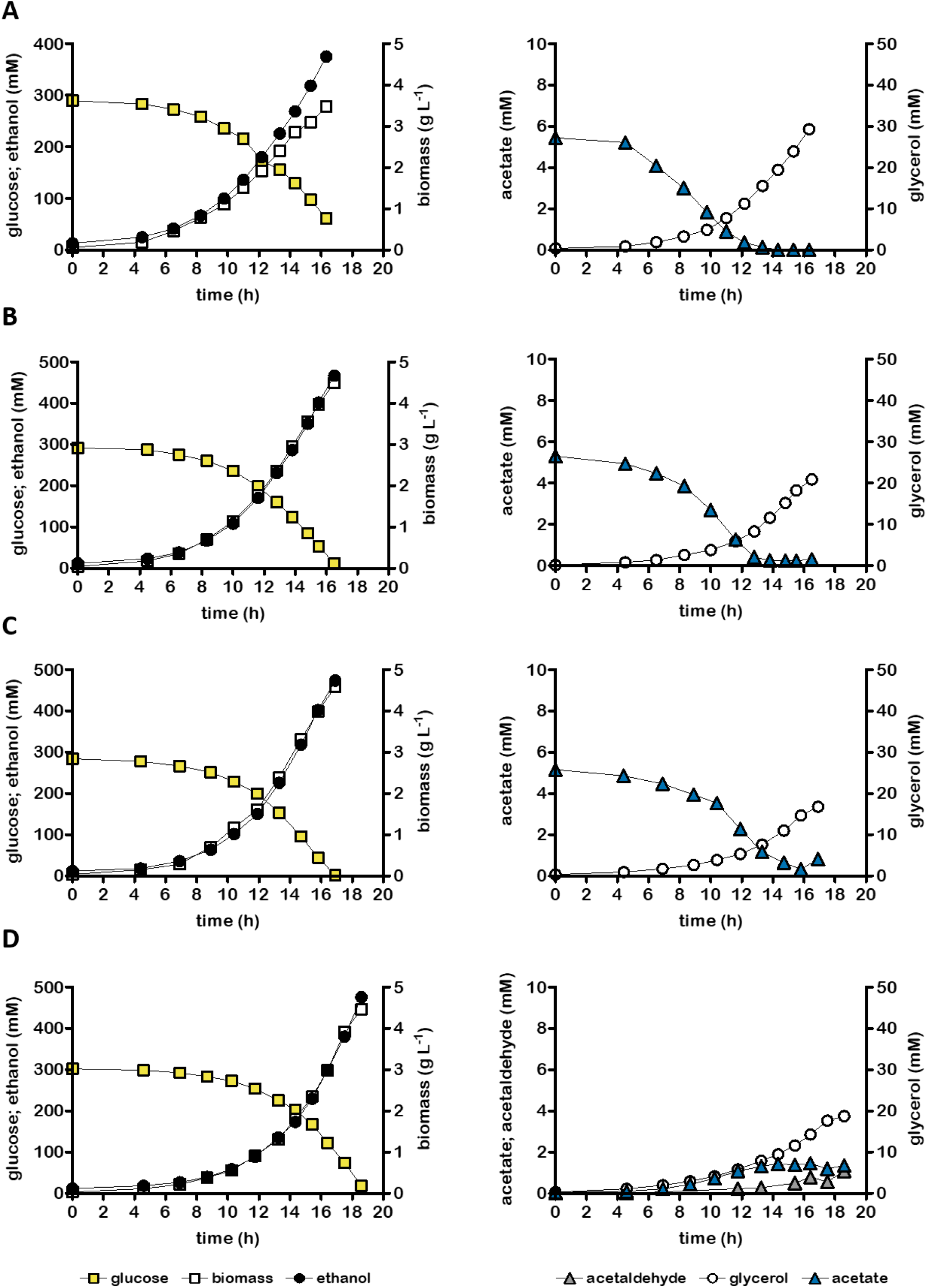
Growth, glucose consumption and product formation of anaerobic bioreactor batch cultures of *S. cerevisiae* strain IMX2503 (*Δgpd2 Δald6 eutE*) (A) and co-cultures of IMX2503 and IMX2736 *(Δgpd2* non-ox PPP↑ *prk* 2× *cbbm groES/groEL*). Non-ox PPP↑ indicates integration of the overexpression cassettes for *RPE1*, *TKL1*, *TAL1*, *NQM1*, *RKI1* and *TKL2*. Cultures were grown on synthetic medium containing 50 g L^-1^ glucose (panel D) or 50 g L^-1^ glucose and 5 mmol L^-1^ acetate (panels A-C) and were inoculated at a ratio of 1.42±0.19 (B), 0.80±0.11 (C) or 1.09±0.02 (D) (inoculum ratio was estimated based on whole genome sequencing). Representative cultures of independent duplicate experiments are shown, corresponding replicate of each culture shown in Fig. S3.

### 2.3 Anaerobic co-cultivation of A-ALD-based and RuBisCO-based strains on glucose

In anaerobic batch cultures grown on glucose, the PRK-RuBisCO-based strain IMX2736 produced acetaldehyde and acetate as byproducts (Table 1, Fig. 2). To investigate whether co-cultivation with an A-ALD-based strain could reduce or eliminate this undesirable byproduct formation, anaerobic batch cultures of strains IMX2736 and IMX2503 were grown on glucose (50 g L^-1^) as sole carbon source. In these cultures, which were grown with an inoculum ratio of 1.1:1 (IMX2503:IMX2736), strain IMX2503 (*Δgpd2 Δald6 eutE*) can only use acetate and acetaldehyde generated by strain IMX2736. Yields of acetate and acetaldehyde were 84% and 72% lower, respectively, than in corresponding monocultures of the RuBisCO-based strain IMX2736 (Tables 1 and 2). Co-cultivation did not extend the fermentation time relative to monocultures of strain IMX2736, while the ethanol yield of the consortium was 1.5% higher than that of monocultures of strain IMX2736 and 8.8% higher than that of monocultures of the reference strain IME324 (p = 0.026 and p = 0.034, respectively, Fig. 2, Fig. 4, Tables 1 and 2). The glycerol yield of these co-cultures was 58% lower than that of monocultures of the reference strain IME324, but 55% higher than that of monocultures of the PRK-RuBisCO-containing strain IMX2736 (Fig. 2, Fig. 4, Tables 1 and 2).

In contrast to co-cultures of strains IMX2736 and IMX2503 on 50 g L^-1^ glucose and 5 mmol L^-1^ acetate, the co-culture on 50 g L^-1^ glucose still produced some acetate and acetaldehyde (Table 2, Fig. 3). A dependency of the A-ALD strain IMX2503 on acetate for fast growth (4, 22) was reflected by a decrease of its relative abundance in the mixed culture during the first phase of batch cultivation on 50 g L^-1^ glucose (Fig. 3). Such a decrease was not observed in co-cultures of these strains that, with a similar inoculum ratio of the two strains, were grown on 50 g L^-1^ glucose and 5 mmol L^-1^ acetate (Fig. 3).

**Table 2:**
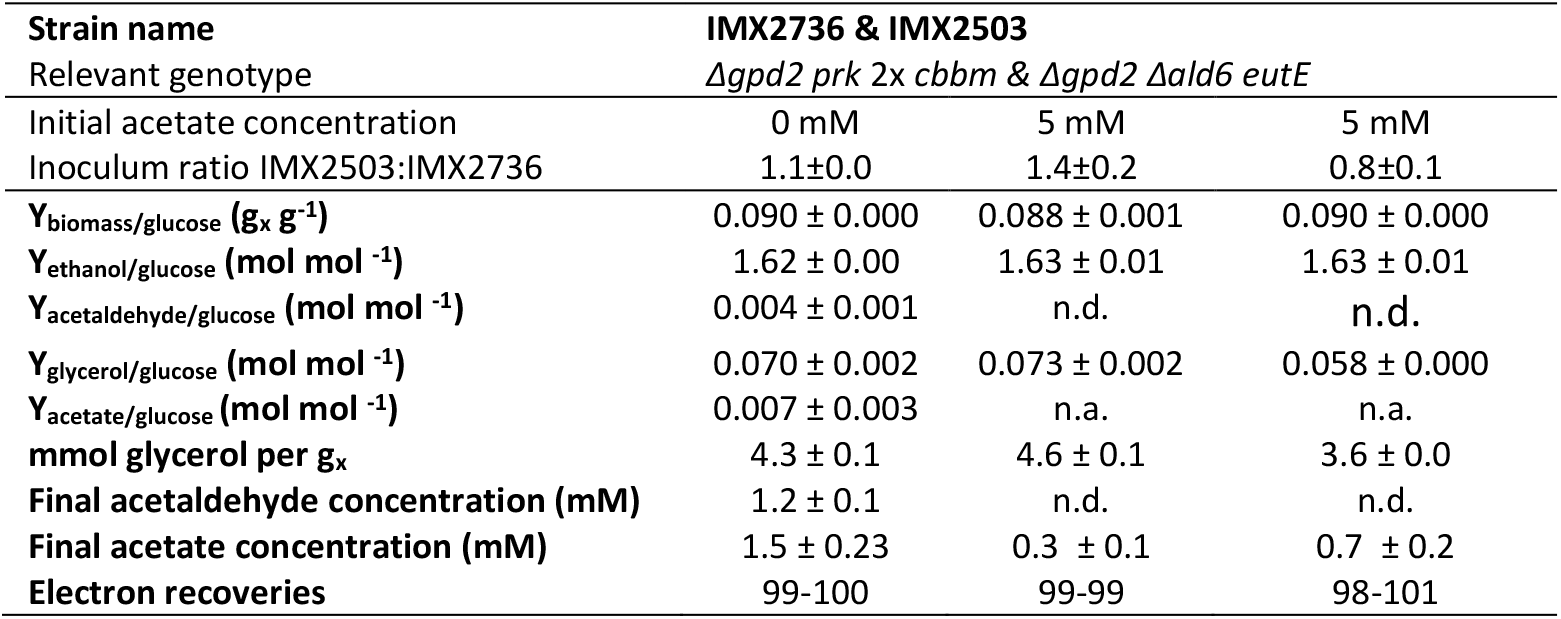
Key physiological parameters of anaerobic bioreactor batch co-cultures of *S. cerevisiae* strains IMX2736 (*Δgpd2* non-ox PPP↑ *prk* 2× *cbbm groES/groEL*) and IMX2503 (*Δgpd2 Δald6 eutE*). Non-ox PPP↑ indicates integration of the overexpression cassettes for *RPE1, TKL1, TAL1, NQM1, RKI1 and TKL2*. Inoculum ratios of the two strains were calculated by genome sequencing. Cultures were grown on synthetic medium with 50 g L-1 glucose, with or without addition of 5 mM acetate. Y indicates yield, subscript x denotes biomass. Acetate and acetaldehyde concentrations indicate values in the culture broth, measured at the end of the cultivation experiments. Yields were calculated using the average of the first two and last two sampling points. Degree-of-reduction balances (61) were used to verify data consistency. Values represent averages ± mean deviations of measurements on independent duplicate cultures for each combination of strain and medium. n.d., not determined; n.a., not applicable (acetate consumption instead of acetate production)

### 2.4. Additional deletion of GPD1 in A-ALD-based strain prevents glycerol production in the absence of acetate

The A-ALD expressing strain IMX2503 (*Δgpd2 Δald6 eutE*) retains a functional *GPD1* gene and therefore, albeit slower than the reference strain (4, 22) can grow in the absence of acetate. Therefore, when the inoculum of co-cultures with the RuBisCO strain IMX2736 contained a high fraction of strain IMX2503 and acetate was consumed before glucose was exhausted, the co-culture displayed a higher glycerol yield (Fig. 4; Table 2). Ideally, depletion of acetate should not lead to enhanced glycerol formation, since this goes at the expense of ethanol yield. Avoiding this requires that the A-ALD strain stops growing when acetate is depleted. We therefore constructed the A-ALD strain IMX2744 (*Δgpd1 Δgpd2 Δald6 eutE*) whose anaerobic growth, due to the elimination of both glycerol-3-phosphate-dehydrogenase isoenzymes, depended on external supply of acetate or acetaldehyde.

As reported for previously constructed *Δgpd1 Δgpd2 EutE* strains (21), strain IMX2744 showed a suboptimal specific growth rate in anaerobic cultures grown on glucose and acetate (Fig. S4). A faster-growing single-cell isolate, IMS1247, was obtained by adaptive laboratory evolution of strain IMX2744 (initial specific growth rate of ca. 0.20 h^-1^) in sequential batch reactors on 50 g glucose L^-1^ and 17 mmol L^-1^ acetate (Fig. S4).

In anaerobic batch cultures on 50 g L^-1^ glucose and 5 mmol L^-1^ acetate, the evolved strain IMS1247 still grew slower than strain IMX2503 (*Δgpd2 Δald6 eutE*) (Fig. 5A and 4A, initial specific growth rates ca. 0.32 h^-1^ and 0.27 h^-1^, respectively). After 19 h, when acetate had been completely consumed, anaerobic cultures of strain IMS1247 had only consumed a quarter of the glucose initially present in the culture (Fig. 5A). However, in contrast to strain IMX2503, strain IMS1247 did not produce any glycerol (Fig. 4A, Fig. 5A).

Whole-genome sequencing of the evolved strain IMS1247 revealed single-nucleotide mutations in *HXK2*, *GIS3* and in the intergenic region in front of *GUT1* (Table S1). The SNP on Chr. 7 location 30770 (*HXK2*) in IMS1247 was used in addition to the two previously described unique SNPs in strain IMX2736. These mutations were used to estimate the ratio of strains IMS1247 and IMX2736 in co-cultures (Fig. 6).

To compensate for the slow growth of strain IMS1247, initial co-cultivation experiments with strain IMX2736 on 50 g L^-1^ glucose and 5 mmol L^-1^ acetate were grown with an inoculum ratio of 5.5:1 (IMS1247:IMX2736). In these cultures, acetate was completely consumed after 16 h, when only half of the glucose had been consumed (Fig. 5B). Complete consumption of glucose occurred after 20 h, which was 3 h later than in monocultures of strain IMX2736 on the same medium (Fig. 2D). This slower conversion was anticipated due to the lower inoculum density of strain IMX2736 and the dependency of strain IMS1247 on exogenous acetate or acetaldehyde.

When the inoculum ratio of the two strains was changed to 1:1 (IMS1247:IMX2736), approximately 1 mmol L^-1^ acetate was left in the culture when glucose was exhausted (Fig. 5C). As a consequence, growth arrest of IMS1247 was prevented and the overall fermentation time was close to that of monocultures of PRK-RuBisCO strain IMX2736 on the same medium (Fig. 6). Glycerol yields in co-cultures of strains IMS1247 and IMX2736 grown on acetate-supplemented medium at inoculum ratios of 5.5 and 1.0 were 77% and 82% lower, respectively, than in corresponding monocultures of the reference strain IME324 and 22% and 37% lower than the monocultures of PRK-RuBisCO strain IMX2736. Moreover, ethanol yields were 8.3% and 7.1% higher, respectively, for co-cultures grown at inoculum ratios of 5.5 and 1.0, than for monocultures of the reference strain IME324 (p = 0.002 and p = 0.014, respectively, Tables 1 and 3).

During anaerobic co-cultivation on 50 g L^-1^ glucose as sole carbon source, growth of the glycerol-negative A-ALD strain IMS1247 was anticipated to depend on supply of acetate and acetaldehyde by the PRK-RuBisCO-based strain IMX2736. Acetate and acetaldehyde yields of the co-cultures of strains IMS1247 and the PRK-RuBisCO IMX2736 on 50 g L^-1^ glucose, inoculated at a ratio of 1.3:1 (IMS1247:IMX2736), were 47% and 61% lower, respectively, than those of a monocultures of strain IMX2736 on 50 g L^-1^ and their ethanol yield was 2.7% higher (p = 0.046, Tables 1 and 3). These byproduct yields were 2.3- and 1.7-fold higher, respectively than observed in co-cultures of strain IMX2736 and strain IMX2503 (*Δgpd2 Δald6 eutE*). The latter observation probably reflects the population dynamics of the co-cultures of strains IMS1247 and IMX2736 on glucose as sole carbon source, which showed a 50% decrease of the relative abundance of strain IMS1247 during fermentation. In contrast, in co-cultures on 50 g L^-1^ glucose and 5 mmol L^-1^ acetate of the two strains with an inoculation ratio of 1, strain IMS1247 represented approximately half of the population throughout fermentation (Fig. 6).

**Fig. 5.**
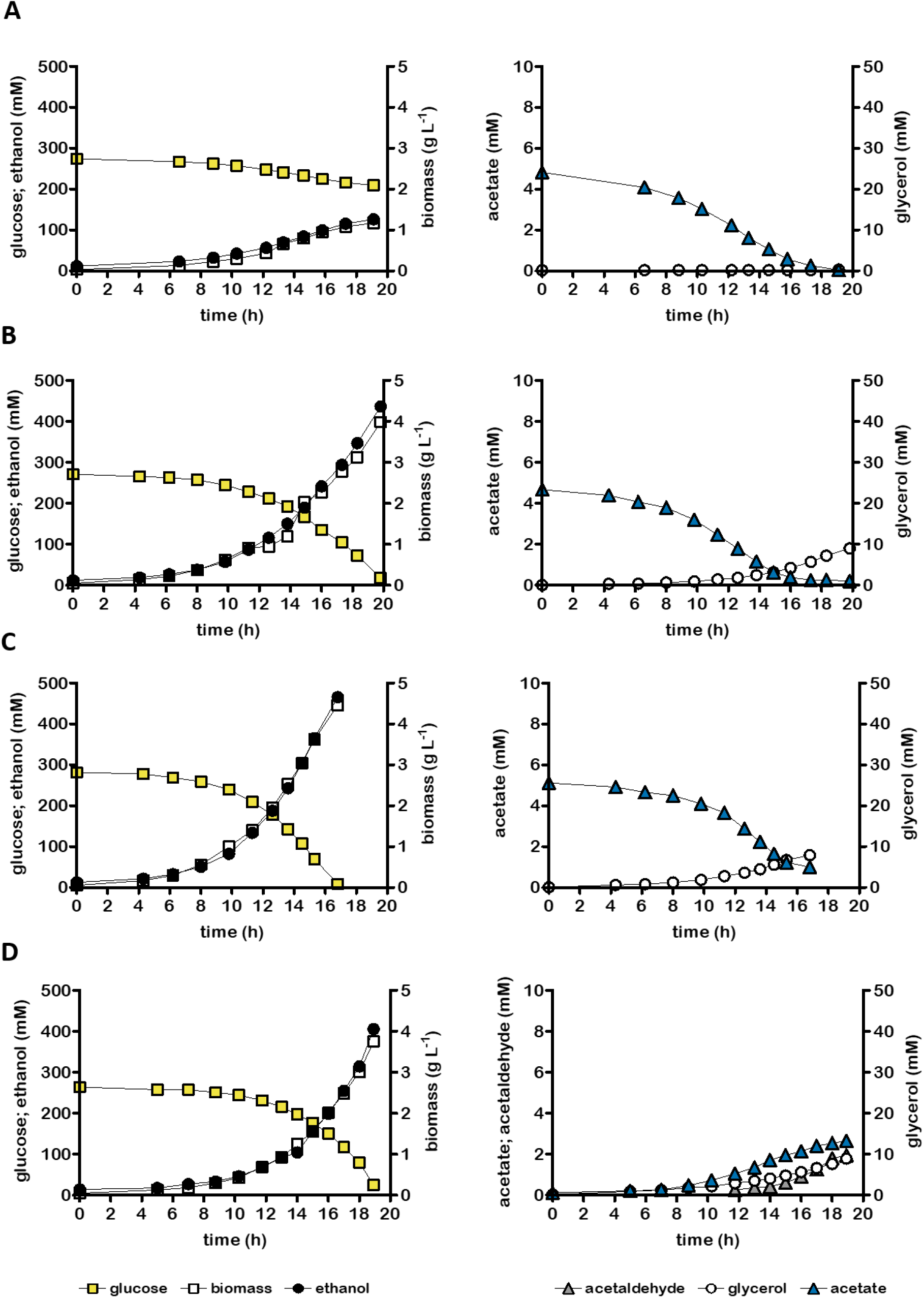
Growth, glucose consumption and product formation of anaerobic bioreactor batch cultures of *S. cerevisiae* strain IMS1247 (*Δgpd1 Δgpd2 Δald6 eutE*, evolved) (**A**) and co-cultures of IMS1247 and IMX2736 (*Δgpd2* non-ox PPP↑ *prk* 2× *cbbm groES*/*groEL*). Non-ox PPP↑ indicates integration of overexpression cassettes for *RPE1*, *TKL1*, *TAL1*, *NQM1*, *RKI1* and *TKL2*. Cultures were grown on synthetic medium containing 50 g L^-1^ glucose (panel **D**) or 50 g L^-1^ glucose and 5 mmol L^-1^ acetate (panels **A**-**C**) and were inoculated at a ratio of 5.5±1.3 (**B**), 1.0±0.2 (**C**) or 1.3±0.4 (**D**). Representative cultures of independent duplicate experiments are shown, corresponding replicate of each culture shown in Fig. S4.

**Fig.6.**
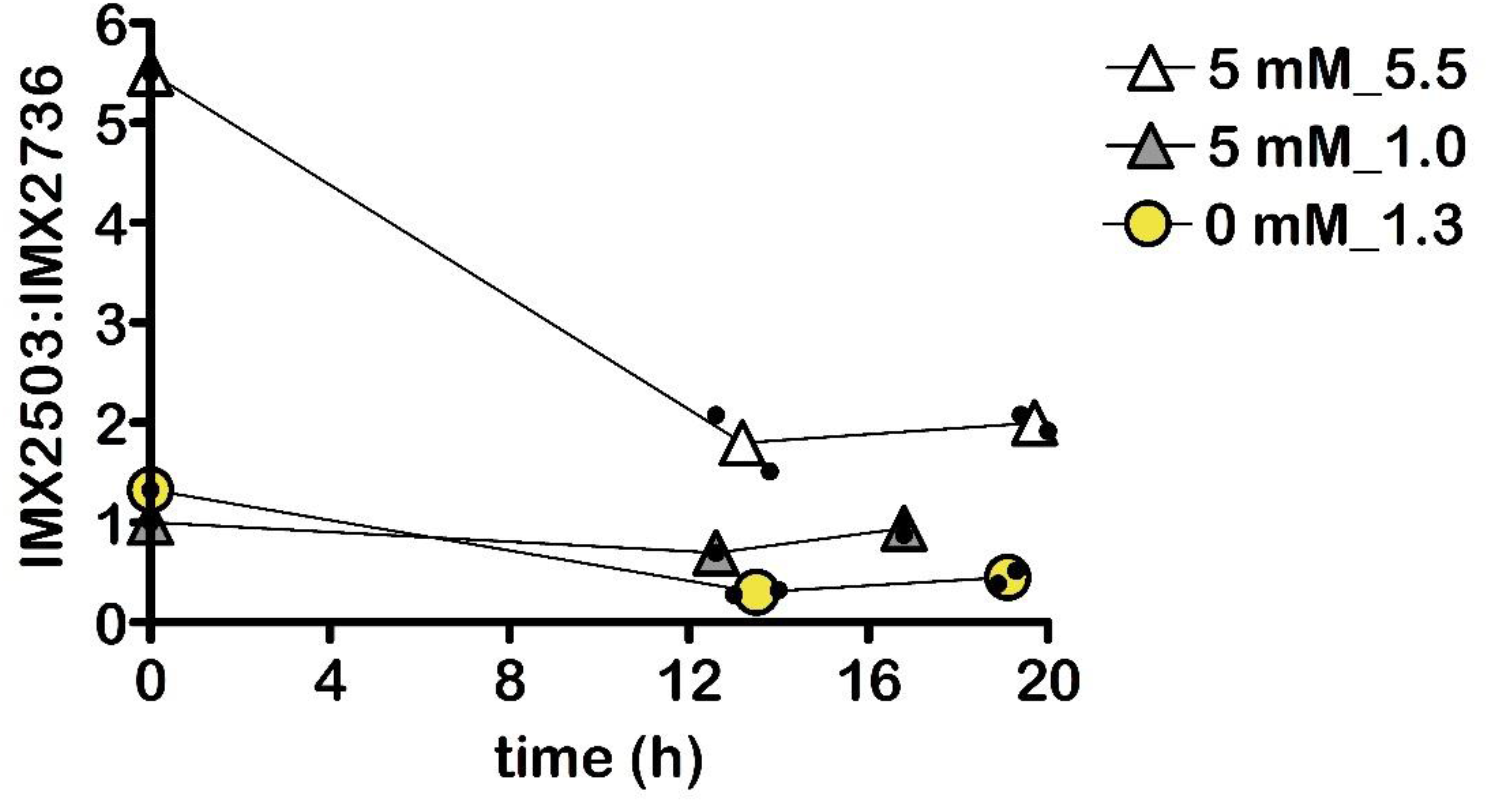
Ratio of *S. cerevisiae* IMS1247 *(Δgpd1 Δgpd2 Δald6 eutE*, evolved) relative to strain IMX2736 *(Δgpd2* non-ox PPP↑ *prk* 2× *cbbm groES/groEL*) in anaerobic bioreactor batch co-cultures on synthetic medium containing 50 g L^-1^ glucose with or without the addition of 5 mM acetate. Ratio was calculated based on whole genome sequencing. Non-ox PPP↑ indicates the integration of the overexpression cassettes for *RPE1*, *TKL1*, *TAL1*, *NQM1*, *RKI1* and *TKL2*. Values represent means and individual values of measurements on independent batch duplicate cultures. Cultures on 50 g L^-1^ glucose and 5 mM acetate were inoculated at two different relative ratios.

**Table 3:**
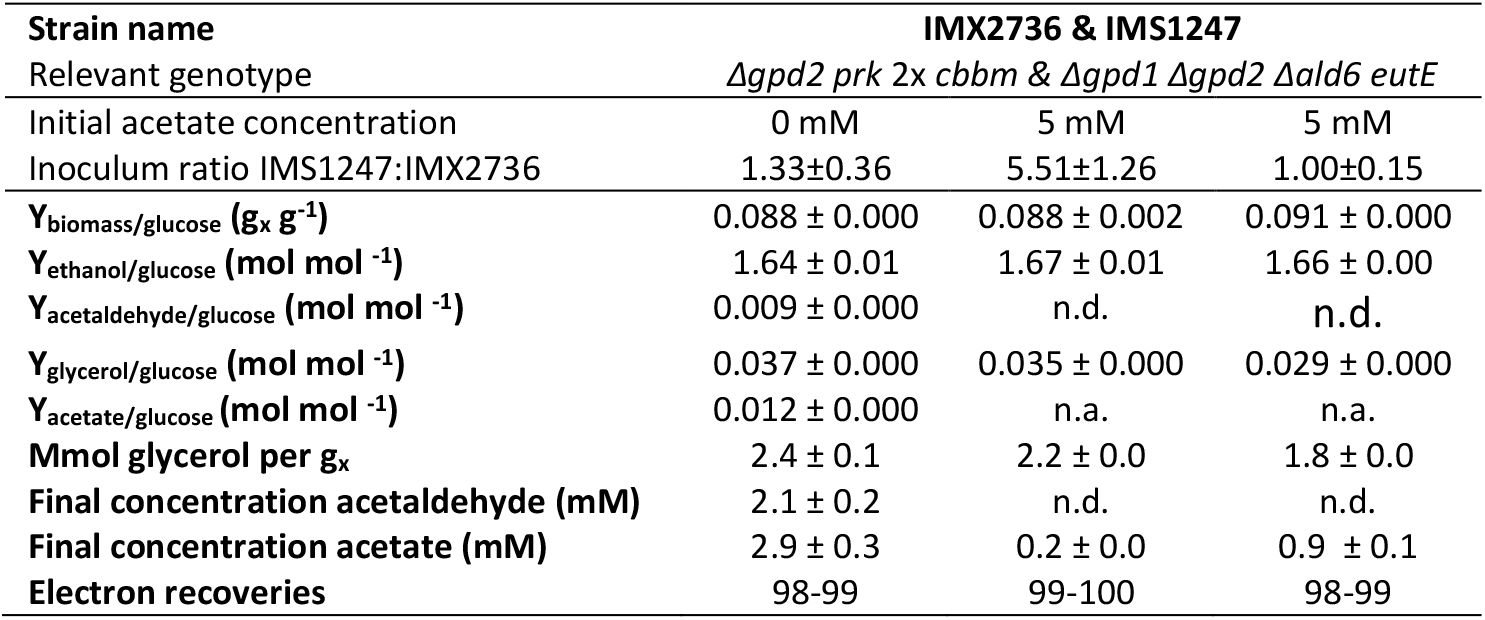
Key physiological parameters of anaerobic bioreactor batch co-cultures of *S. cerevisiae* strains IMX2736 (*Δgpd2*, non-ox PPP↑ *prk* 2× *cbbm groES/groEL*) and IMS1247 (*Δgpd1 Δgpd2 Δald6 eutE*, evolved). Non-ox PPP↑ indicates integration of the overexpression cassettes for *RPE1, TKL1, TAL1, NQM1, RKI1 and TKL2*. Inoculum ratios of the two strains were calculated by genome sequencing. Cultures were grown on synthetic medium with 50 g L^-1^ glucose, with or without addition of 5 mM acetate. Y indicates yield, subscript x denotes biomass. Acetate and acetaldehyde concentrations indicate values in the culture broth, measured at the end of the cultivation experiments. Yields were calculated using the average of the first two and last two sampling points. Degree-of-reduction balances (61) were used to verify data consistency. Values represent averages ± mean deviations of measurements on independent duplicate cultures for each combination of strain and medium. n.d., not determined; n.a., not applicable (acetate consumption instead of acetate production)

## 3. Discussion

Advantages of microbial interactions are well described for multi-species natural microbial ecosystems and microbial consortia applied in food fermentation processes (36, 37). Previous laboratory studies on the use of consortia of different engineered *S. cerevisiae* strains for ethanol production focused on conversion of polysaccharides or sugar mixtures by consortia of engineered ‘specialist’ strains (38, 39). The present study focused on efficient conversion of glucose to ethanol, both in the absence and presence of low concentrations of acetate in growth media. Its results show that consortia of engineered PRK-RuBisCO-based and A-ALD-based *S. cerevisiae* strains enable higher ethanol yields than obtained with a non-engineered reference strain, while reducing net formation of byproducts originating from PRK-RuBisCO-based *S. cerevisiae* and removing acetate from growth media.

Introduction of PRK and RuBisCO, combined with modifications in the central metabolism of *S. cerevisiae*, enables improved ethanol yields in fast-growing anaerobic cultures (15). However, during slower anaerobic growth in continuous cultures (22), and in batch cultures on 50 g L^-1^ glucose (Fig. 2 CD), PRK-RuBisCO strains optimized for fast growth generate acetaldehyde and acetate as byproducts. In a recent study, PRK-RuBisCO strains into which an A-ALD pathway had been introduced showed inferior acetate reduction relative to a strain that only expressed the A-ALD pathway (22). A lack of *in vivo* reductive A-ALD activity in ‘dual pathway’ strains was attributed to the impact of acetaldehyde and a low NADH/NAD^+^ ratio, both generated by activity of the PRK-RuBisCO bypass, on the reversible A-ALD reaction (ΔG^0^’ = 17.6 kJ mol^-1^ for the reductive reaction, (40)). Further engineering to adapt *in vivo* activity of PRK and RuBisCO in dual-pathway strains to NADH availability in dynamic industrial cultures would require introduction of dynamic regulation circuits (22). Alternatively, our results show that interference of the two pathways was mitigated by their compartmentation in separate co-cultivated strains. This co-cultivation approach enabled high ethanol yields while strongly reducing acetate and acetaldehyde production in glucose-grown batch cultures (Fig. 4D, Fig 5D). Minimizing acetaldehyde production is not only relevant for improving ethanol yield but also to prevent its toxicity to yeast cells (28, 29) and also in view of environmental and health issues (41, 42).

Consortia of PRK-RuBisCO and A-ALD-based strains removed essentially all acetate from media with acetate-to-glucose ratios similar to those in feedstocks for first-generation bioethanol production (Fig. 4 BC, Fig. 5 BC). The complete removal of acetate during fermentation does not only contribute to increased ethanol yields, but also prevents its recycling into subsequent fermentation runs and, potentially its accumulation to inhibitory levels, as a result of recycling thin stillage and evaporator condensate (43).

Co-cultivation of the PRK-RuBisCO strain IMX2736 with the A-ALD strains IMX2503 (*Δgpd2 Δald6 eutE*) and IMS1247 (*Δgpd1 Δgpd2 Δald6 eutE*) revealed a trade-off between stoichiometry and kinetics (Fig. 4, Fig. 5). Co-cultivation with strain IMS1247, which could not produce any glycerol, led to low glycerol formation but led to extended fermentation times (Fig. 5 B). Ideally, low-glycerol growth and vigorous fermentation by A-ALD-based strains should continue even when acetate availability in the medium becomes growth limiting. In addition to the option of supplying small amounts of acetate during fermentation, this goal may be pursued by further strain engineering to allow for a strongly constrained rate of glycerol production. Alternatively, *S. cerevisiae* strains may be used in which A-ALD reduces acetyl-CoA synthesized from glucose via engineered pathways that generates fewer than 2 mole of NADH per mole of acetyl-CoA. Such strategies can be based on introduction of a heterologous pyruvate-formate lyase (PFL) (44, 45), or of a heterologous phosphoketolase (PK) and phosphotransacetylase (PTA) (46–48).

With the exception of classical processes in Brazil, in which yeast biomass is recycled after each fermentation run (49, 50), industrial batch fermentation processes for ethanol production are typically started with fresh pre-cultures of yeast strains provided by specialist companies. When exclusively relying on metabolic engineering of monocultures, adaptations to regular changes in feedstock and process configurations would require extensive strain engineering. Use of co-cultures enables fast adaptation of inoculation ratios of available strains to optimally balance ethanol yield, productivity and byproduct formation (Fig. 4, Fig. 5). The readiness of the first-generation bioethanol industry to adopt this strategy is illustrated by a recent report that, without disclosing details on the strains involved, indicates that blends of engineered yeast strains are already applied at full industrial scale (51).

## 4. Materials and Methods

### 4.1. Strains, media and maintenance

The *S. cerevisiae* strains used in this study (Table 4) were derived from the CEN.PK lineage (52, 53). Synthetic medium (SM), containing 3.0 g L^-1^ KH2PO4, 0.5 g L^-1^ MgSO4·7H2O, 5.0 g L^-1^ (NH4)2SO4, trace elements and vitamins, was prepared as described previously (54). Shake-flask cultures were grown on SM supplemented with 20 g L^-1^ glucose (SMD) and bioreactor cultures on SM with 50 g L^-1^ glucose. Anaerobic growth media were supplemented with ergosterol (10 mg L^-1^) and Tween 80 (420 mg L^-1^) (55). Anaerobic cultures were grown on SMD in which the KH2PO4 concentration was raised to 14.4 g L^-1^ for extra pH buffering. Complex medium (YPD) contained 10 g L^-1^ Bacto yeast extract (Thermo Fisher Scientific, Waltham MA), 20 g L^-1^ Bacto peptone (Thermo Fisher Scientific) and 20 g L^-1^ glucose. To select for presence of an acetamidase marker cassette (56), (NH4)2SO4 was replaced by 6.6 g L^-1^ K2SO4 and 0.6 g L^-1^ filter-sterilized acetamide. Where indicated, pure acetic acid solution (≥99.8%, Honeywell, Charlotte NC) was added to media at a concentration of 0.30 g L^-1^ or 1.0 g L^-1^. *Escherichia coli* XL1-Blue cultures were grown on lysogeny broth (LB) (57) containing 10 g L^-1^ tryptone (Brunschwig Chemie B.V., Amsterdam, The Netherlands), 5.0 g L^-1^ yeast extract and 10 g L^-1^ NaCl. Where relevant, LB was supplemented with 100 mg L^-1^ ampicillin (Merck, Darmstadt, Germany). *E. coli* strains were grown overnight at 37 °C in 15-mL tubes containing 5 mL LB, shaken at 200 rpm in an Innova 4000 Incubator (Eppendorf AG, Hamburg, Germany). Solid media were prepared by adding 20 g L^-1^ agar (Becton Dickinson, Breda, The Netherlands) prior to heat sterilization. *S. cerevisiae* plate cultures were incubated at 30 °C until colonies appeared (1-5 d), while *E. coli* plates were incubated overnight at 37 °C. Frozen stock cultures were prepared by freezing samples from fully grown batch cultures at -80 °C after addition of 30% (v/v) glycerol.

**Table 4.** *S. cerevisiae* strains used in this study. *Kl* denotes *Kluyveromyces lactis*.

| Strain name | Relevant genotype | Parental strain | origin |
| --- | --- | --- | --- |
| CEN.PK113-5D | *MATa ura3-52* | - | (52) |
| IMX581 | *MATa ura3-52 can1::cas9 natNT2* | CEN.PK113-5D | (58) |
| IME324 | *MATa ura3-52 can1::cas9 natNT2 p426-TEF(empty)* | IMX581 | (15) |
| IMX2503 | *MATa ura3-52 can1::cas9 natNT2 ALD6::pTDH3-eutE gpd2Δ pUDR774 (KIURA3)* | IMX581 | (22) |
| IMX2744 | *MATa ura3-52 can1::cas9 natNT2 ALD6::pTDH3-eutE gpd2Δ gpd1Δ pUDR774 (KIURA3)* | IMX2503 | This study |
| IMS1247 | Strain IMX2744 evolved for faster anaerobic growth on 50 g L<sup>-1</sup> glucose and 1 g L<sup>-1</sup> acetic acid | IMX2744 | This study |
| IMX2736 | *MATa ura3-52 can1::cas9 natNT2 gpd2::(pTDH3-RPE1, pPGK1-TKL1, pTEF1-TAL1, pPGI1-NQM1, pTPI1-RKI1, pPYK1-TKL2) sga1::(pDAN1-prk, pTDH3-cbbm (2 copies) pTPI1-groES, pTEF1-groEL) pUDR103 (KIURA3)* | - | (17) |

### 4.2. Plasmid construction

Cas9 target sequences in *GPD1* and *GPD2* were identified as described previously (58). To construct plasmid pUDR203, a linear backbone fragment of pROS11 was first PCR amplified with primer 5793. Subsequently, DNA fragments encoding *GPD1*-targeting and *GPD2*-targeting gRNA cassettes were PCR-amplified using primers 6965/6966. Phusion high-fidelity DNA Polymerase (Thermo Fisher) was used as specified by the manufacturer. Plasmid-backbone and insert fragments were isolated from gels with the Zymoclean Gel DNA Recovery kit (Zymo Research, Irvine CA). DNA concentrations were measured with a NanoDrop 2000 spectrophotometer (Thermo Fisher) at a wavelength of 260 nm. Plasmid assembly was performed by in vitro Gibson Assembly using a HiFi DNA Assembly master mix (New England Biolabs, Ipswich, MA), downscaled to 5 µL reaction volumes. 1 µL of the reaction mixture was used to transform *E. coli* XL-1 Blue cells with a heat-shock protocol (59). Plasmid pUDR203 was isolated from *E. coli* XL-I Blue cells with the Sigma GenElute Plasmid Miniprep Kit (Sigma-Aldrich) as specified by the manufacturer.

**Table 5.** Plasmids used in this study

| Plasmid | Characteristics | Reference |
| --- | --- | --- |
| pROS11 | 2 $\mu\text{m}$ ori, *AmdS*, gRNA-*CAN1.Y* gRNA-*ADE2.Y* | (58) |
| pUDR103 | 2 $\mu\text{m}$ ori, *KIURA3*, gRNA-*SGA1.Y* | (15) |
| pUDR203 | 2 $\mu\text{m}$ ori, *AmdS*, gRNA-*GPD1.Y* gRNA-*GPD2.Y* | This work |
| pUDR774 | 2 $\mu\text{m}$ ori, *KIURA3*, gRNA-*GPD2.Y* gRNA-*GPD2.Y* | (22) |

### 4.3. Genome editing

A dsDNA-repair fragment for deletion of *GDP1* was obtained by mixing primers 6969/6970 in a 1:1 molar ratio. This mixture was heated to 95 °C for 5 min and allowed to cool to room temperature. *S. cerevisiae* IMX2744 was constructed by co-transforming strain IMX2503 with gRNA-plasmid pUDR203 and the *GPD1* repair fragment, using the lithium-acetate method (60). Transformants were selected on SM supplemented with 20 g L^-1^ glucose (SMD) and acetamide (56), after which correct deletion of *GPD1* was checked by diagnostic colony PCR with DreamTaq polymerase (Thermo Fisher). pUDR203 was removed by growing non-selectively on SMD, while pUDR774 was retained to support uracil prototrophy. A correct transformant was restreaked thrice on SMD and stored at -80 °C.

### 4.4. Anaerobic shake-flask cultivation

Anaerobic shake-flask cultures of single-colony isolates from SBR evolution experiments with *S. cerevisiae* IMX2744 were grown at 30 °C in 50-mL round-bottom shake-flasks containing 30 mL extra buffered SM supplemented with vitamins, 50 g L^-1^ glucose, 1 g L^-1^ acetic acid and Tween 80/ergosterol. Seven single-colony isolates from each reactor isolated after 38 repeated-batch cycles, along with four isolates from reactor I and seven from reactor II isolated after 63 cycles, were analysed for their specific growth rates. Of these 25 single colony isolates, IMS1247 was selected with the fastest growth rate (0.27 h^-1^), which was isolated from reactor I after 63 cycles. Shake flasks were placed on an IKA KS 260 basic shaker (Dijkstra Verenigde BV, Lelystad, The Netherlands, 200 rpm) in a Bactron anaerobic chamber (Sheldon Manufacturing Inc., Cornelius, OR) under an atmosphere of 5% (v/v) H2, 6% (v/v) CO2 and 89% (v/v) N2 (55).

### 4.5. Bioreactor cultivation

Anaerobic bioreactor batch and sequential-batch cultures were grown at 30 °C in 2-L bioreactors (Applikon, Delft, The Netherlands). Culture pH was maintained at 5.0 by automatic addition of 2 M KOH. Bioreactor cultures were grown on SM, supplemented with glucose (50 g L^-1^), acetic acid (0.3 g L^-1^ or 1 g L^-1^ as indicated), Tween 80 (420 mg L^-1^) and ergosterol (10 mg L^-1^), and antifoam C (0.2 g L^-1^) (Sigma-Aldrich). Bioreactor cultures were operated at a working volume of 1 L and sparged at 0.5 L min^-1^ with an N2/CO2 (90/10%) gas mixture, except for the laboratory-evolution cultures of strain IMX2744 and the anaerobic bioreactor batch of strains IMX2744 and IMS1247 on 50 g L^-1^ of glucose and 1 g L^-1^ of acetic acid, which were sparged with pure N2. The outlet gas stream was cooled to 4 °C in a condenser to minimize evaporation. Oxygen diffusion was minimized by use of Norprene tubing (Saint-Gobain, Amsterdam, The Netherlands) and Viton O-rings (ERIKS, Haarlem, The Netherlands) (55). Inocula for bioreactor cultures were prepared in 500-mL shake flasks containing 100 mL SMD. A first preculture, inoculated with a frozen stock culture and grown aerobically at 30 °C for 15-18 h, was used to inoculate a second preculture. Upon reaching mid-exponential phase (OD660 of 3-6), the second preculture was used to inoculate a bioreactor culture at an initial OD660 of 0.2-0.4. For inoculation of the co-culture, the OD660 was used to calculate how much volume from each strain needed to be added into the reactor. DNA isolated from the mix containing both strains, used for inoculation was sequenced to determine the estimated starting inoculum ratio.

Laboratory evolution of *S. cerevisiae* IMX2744 was performed in sequential batch reactor (SBR) set-ups on SM supplemented with 50 g L^-1^ glucose and 1 g L^-1^ acetic acid. Cultures were sparged at 0.5 L min^-1^ with pure N2. When, after having peaked, the CO2 concentration in the off-gas had decreased to 60% of the highest CO2-value measured during the preceding batch cycle, an effluent pump was automatically switched on for 20 min. After this emptying phase only 0.05 L broth remained in the reactor. The effluent pump was then stopped and the inflow pump activated to supply fresh sterile medium from a 20-L reservoir vessel. This refill phase was stopped via an electrical level sensor calibrated at a 1-L working volume.

### 4.6. Analytical methods

The optical density of cultures was measured at 660 nm on a Jenway 7200 spectrophotometer (Bibby Scientific, Staffordshire, UK). Biomass dry weight was measured as described previously (14). Metabolite concentrations were determined by high-performance liquid chromatography and a first-order evaporation rate constant of 0.008 h^-1^ was used to correct ethanol concentrations (14). Acetaldehyde concentrations in culture broth were determined after derivatization with 2,4-dinitrophenylhydrazine as described previously (17). As carbon recoveries could not be accurately calculated due to the high concentration of CO2 in the inlet gas of bioreactor cultures, electron recoveries were used instead (61).

### 4.7. Whole-genome sequencing

Genomic DNA was extracted from an 100-mL aerobic, late-exponential-phase (OD660 of 10– 15) shake-flask culture on SMD of *S. cerevisiae* strain IMX2503, using a Qiagen Blood & Cell Culture DNA kit and 100/G Genomics-tips (Qiagen, Hilden, Germany). Custom paired-end sequencing of genomic DNA was performed by Macrogen (Amsterdam, The Netherlands) on a 350-bp PCR-free insert library using Illumina SBS technology. Sequence reads were mapped against the genome of *S. cerevisiae* CEN.PK113-7D (62) to which a virtual contig containing p*TDH3*-*eutE* had been added, and processed as described previously (63).

To determine relative abundancy of strains IMX2736 and IMX2503 in bioreactor batch co-cultivation experiments, 50-mL culture samples were centrifuged for 10 minutes at 5,000 × *g* and biomass pellets temporarily stored at -20 °C. These pellets were used for genomic DNA extraction and Illumina sequencing. Sequence reads were mapped against the genome of *S. cerevisiae* CEN.PK113-7D to which a virtual contig containing p*TDH3*-*eutE*, p*DAN1*-*PRK*, p*TDH3*-*cbbM*, p*TPI1*-g*roES* and p*TEF1*-*groEL* had been added. Unique SNPs in strains IMX2736 and IMS1247 were identified and used to quantify the percentage of each strain present in the sample.

### 4.9. Statistical analysis

Significance was assessed by performing a two-sided unpaired Student’s t-test. Differences were considered to be significant if a p-value < 0.05 was obtained.

## Supplementary information

**Additional file 1:** Measurement data of anaerobic batch cultures of IME324, IMX2736, co-cultures of IMX2503:IMX2726 and IMS1237:IMX2736 on 50 g L^-1^ of glucose. Data were used to prepare Fig. 2-6 of the manuscript and Fig. S1-S3 and Fig. S5 of the supplementary materials.

**Additional file 2:** Measurement data of anaerobic batch cultures of IMX2736, IMS1247, IMX2744, co-cultures of IMX2503:IMX2726 and IMS1237:IMX2736 on 50 g L^-1^ of glucose and 5 mmol L^-1^ of acetate. Data were used to prepare Figure 2-6 of the manuscript and Fig. S1-S3 and Fig. S5 of the supplementary materials.

**Additional file 3:** Measurement data of anaerobic batch cultures of IMX2744 and IMS1247 on 50 g L^-1^ of glucose and 1 g L^-1^ of acetate. Data were used to prepare Fig. S4 of the supplementary materials.

**Additional file 4:** Evolution using SBR of IMX2744 for faster anaerobic growth on 50 g L^-1^ of glucose and 1 g L^-1^ of acetate. Data were used to prepare Fig. S4 of the supplementary materials.

**Additional file 5:** Supplementary materials. Fig. S1: Specific growth rates at different timepoints of anaerobic batch cultures of strains IME324 and IMX2736. Fig. S2: Growth, glucose consumption, ethanol formation, acetaldehyde formation and glycerol formation in anaerobic bioreactor batch cultures of IME324 and IMX2736. Fig. S3: Growth, glucose consumption, ethanol formation, acetaldehyde formation and glycerol formation in anaerobic bioreactor batch co-cultures of IMX2503 and IMX2736. Fig. S4: Evolution using SBR of IMX2744 for faster anaerobic growth on 50 g L^-1^ of glucose and 1 g L^-1^ of acetate. Fig. S5: Growth, glucose consumption, ethanol formation, acetaldehyde formation and glycerol formation in anaerobic bioreactor batch co-cultures of IMS1247 and IMX2736. Table S1: oligonucleotide primers used in this study.

## Declarations

### Author contributions

**AA:** Validation, Methodology, Formal analysis, Investigation, Writing – Original Draft, Writing – Review & Editing, Visualization. **IM**: Investigation, Writing – Review & Editing. **MJ:** Writing – Review & Editing. **RM:** Conceptualization, Supervision, Writing – Review & Editing. **JP:** Conceptualization, Supervision, Writing – Original Draft, Writing – Review & Editing.

### Data availability

Short read DNA sequencing data of the *Saccharomyces cerevisiae* strain IMS1247 were deposited at NCBI under BioProject accession number PRJNA972873. All measurement data used to prepare Fig. 2, Fig. 3, Fig. 4, Fig. 5, Fig. 6, Table 1, Table 2 and Table 3 of the manuscript and Fig. S1, Fig. S2, Fig. S3, Fig. S4 and Fig. S5 are available in Additional files 1, 2, 3 and 4.

## Acknowledgements

We want to thank Rinke van Tatenhove-Pel for stimulating discussions.

