## Supplementary materials for "Co-cultivation of *Saccharomyces cerevisiae* strains combines advantages of different metabolic engineering strategies for improved ethanol yield"

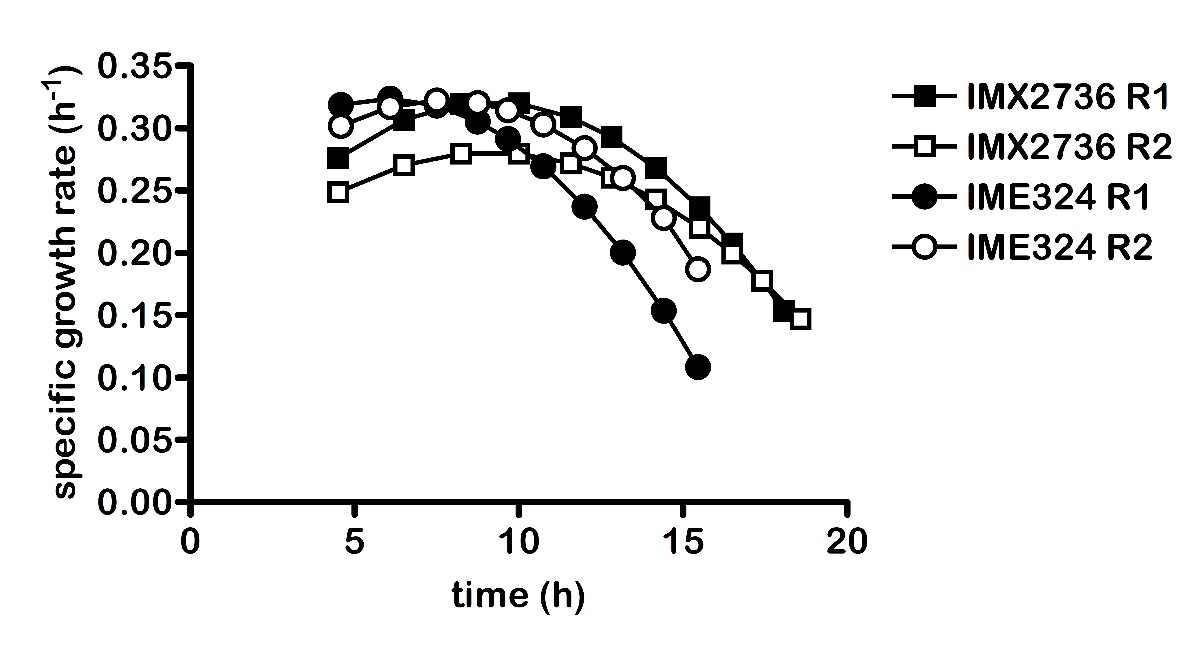

**Figure S1:** Specific growth rates at different timepoints of anaerobic batch cultures of S. cerevisiae strains IME324 (reference) and IMX2736 (non-ox PPP↑ Δgpd2 pDAN1-prk 2x pTDH3-cbbm pTPI1-groES pTEF1-groEL). Cultures were grown on synthetic medium with 50 g L^-1^ of glucose at pH 5 and at 30 °C and sparged with a 90:10 mixture of N_2_ and CO_2_. Non-ox PPP↑ indicates the integration of the expression cassettes of pTDH3-RPE1, pPGK1-TKL1, pTEF1-TAL1, pPGI1-NQM1, pTPI1-RKI1 and pPYK1-TKL2.

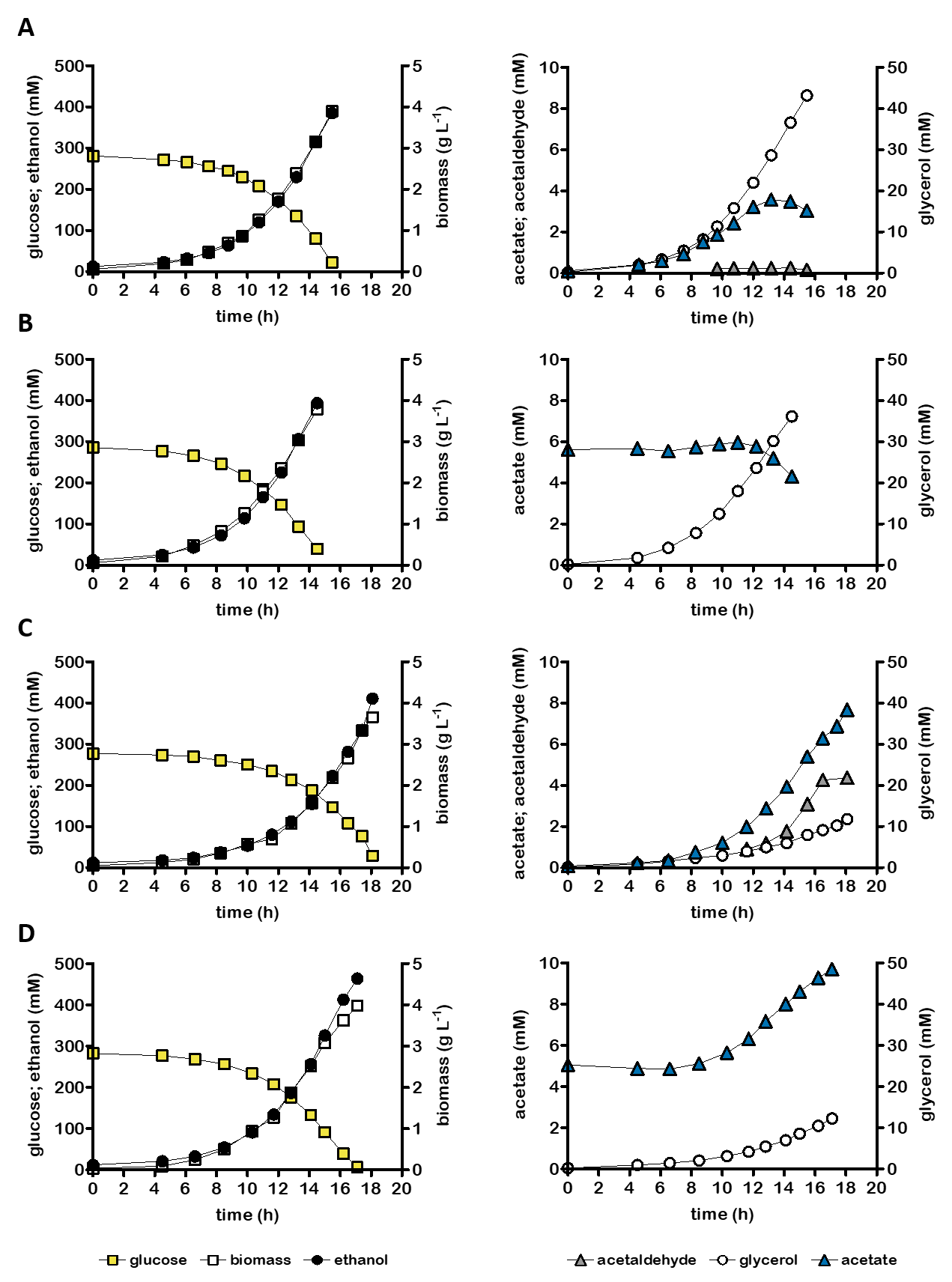

**Figure S2:** Growth, glucose consumption and product formation of anaerobic bioreactor batch cultures of individual S. cerevisiae strains, grown on SM with 50 g L^-1^ glucose (panels **A** and **C**) or on SM with 50 g L^-1^ glucose and 5 mmol L^-1^ acetate (panels **B** and **D**). Panels show data for S. cerevisiae strains IME324 (reference strain, **A** and **B**) and IMX2736 (∆gpd2, non-ox PPP↑, prk, 2x cbbm, groES, groEL, **C** and **D**) Non-ox PPP↑ indicates integration of the overexpression cassettes for RPE1, TKL1, TAL1, NQM1, RKI1 and TKL2. Representative cultures of independent duplicate experiments are shown, corresponding replicate of each culture shown in Figure 1.

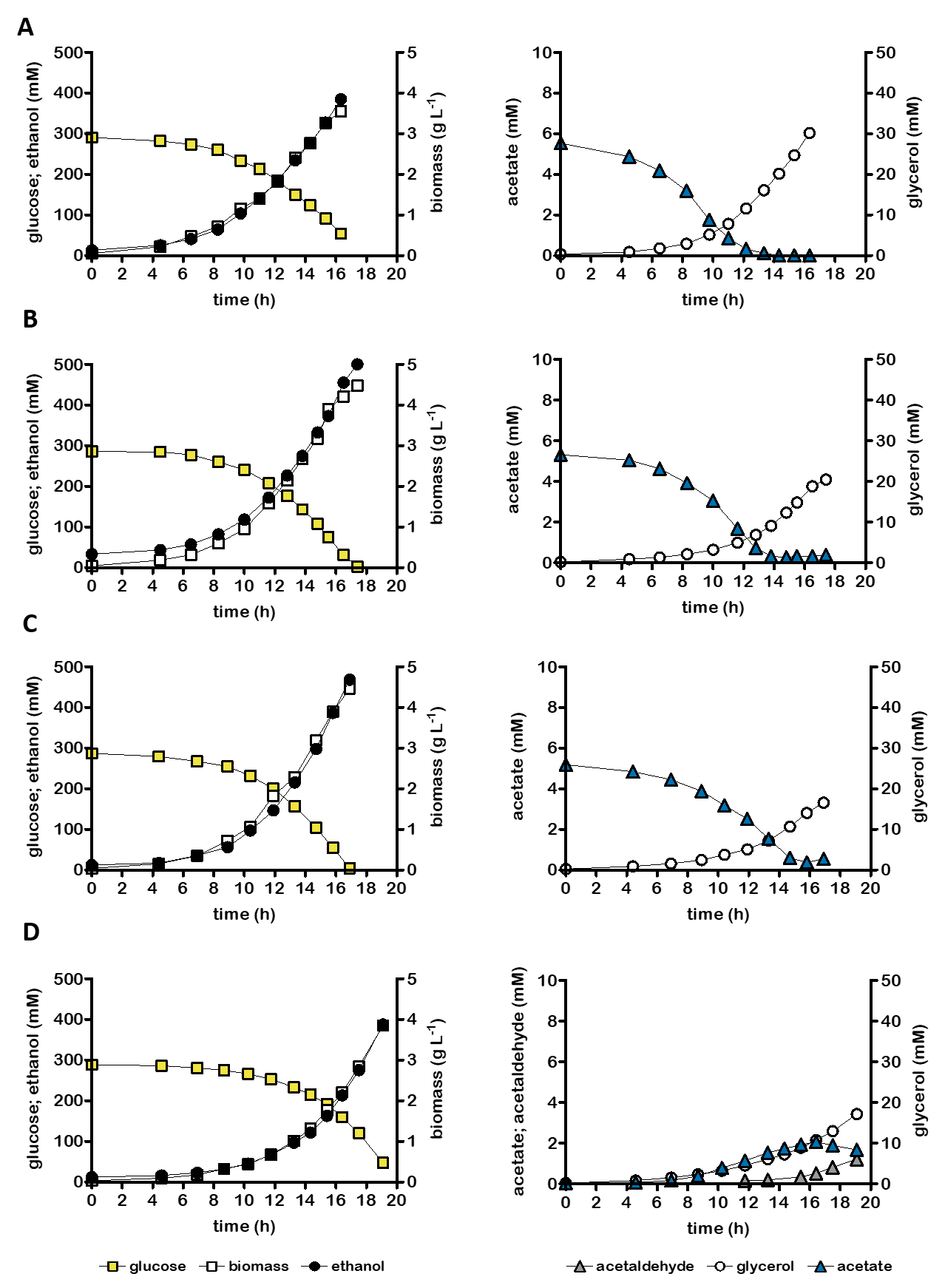

**Figure S3:** Growth, glucose consumption and product formation of anaerobic bioreactor batch cultures of S. cerevisiae strain IMX2503 (∆gpd2 ∆ald6 eutE) (**A**) and co-cultures of IMX2503 and IMX2736 (∆gpd2, non-ox PPP↑, prk, 2x cbbm, groES, groEL). Non-ox PPP↑ indicates integration of the overexpression cassettes for RPE1, TKL1, TAL1, NQM1, RKI1 and TKL2. Cultures were grown on synthetic medium containing 50 g L^-1^ glucose (panel **D**) or 50 g L^-1^ glucose and 5 mmol L^-1^ acetate (panels **A**-**C**) and were inoculated at a ratio of 1.4±0.2 (**B**), 0.8±0.1 (**C**) or 1.1±0.0 (**D**) (inoculum ratio was estimated based on whole genome sequencing). Representative cultures of independent duplicate experiments are shown, corresponding replicate of each culture shown in Figure 3.

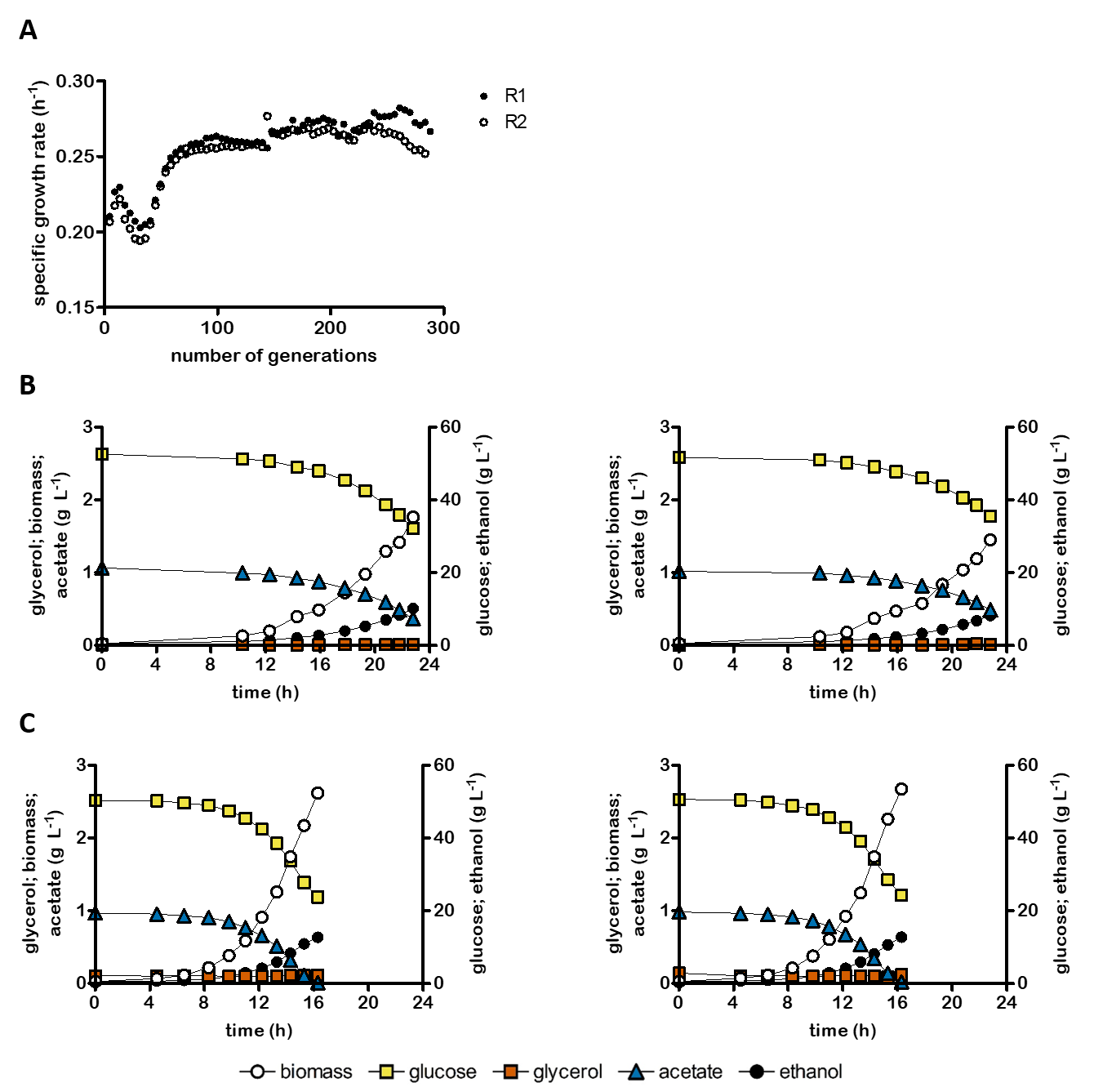

**Figure S4:** evolution performed in SBR set-up to increase the specific growth rate of IMX2744 in anaerobic batch bioreactors on 50 g L^-1^ of glucose and 1 g L^-1^ of acetate. Overview of the specific growth rate as function of the number of generations (**A**). Growth, glucose consumption and product formation of both replicate anaerobic bioreactor batch cultures of S. cerevisiae strain IMX2744 (∆gpd1 ∆gpd2 ∆ald6 eutE) (**B**) and evolved single colony isolate IMS1247 (**C**).

**Table S1:** Identified mutation based on whole genome sequencing in IMS1247 (∆gpd1 ∆gpd2 ∆ald6 eutE) compared to IMX2503 (∆gpd2 ∆ald6 eutE).

| **Position** | **Location annotation** | **Mutation** | **Altered aa** | **Gene annotation** |
| --- | --- | --- | --- | --- |
| Chromosome 7, position 30790 | Coding sequence of *HXK2* | G 🡪 A | G177S | Hexokinase isoenzyme 2 |
| Chromosome 8, position 35455 | Intergenic, in front of *GUT1* | A 🡪 T | n.a. | n.a. |
| Chromosome 8, position 35456 | Intergenic, in front of *GUT1* | T 🡪 A | n.a. | n.a. |
| Chromosome 12, position 307396 | Coding sequence of *GIS3* | C 🡪 T | S288S | Protein of unknown function |

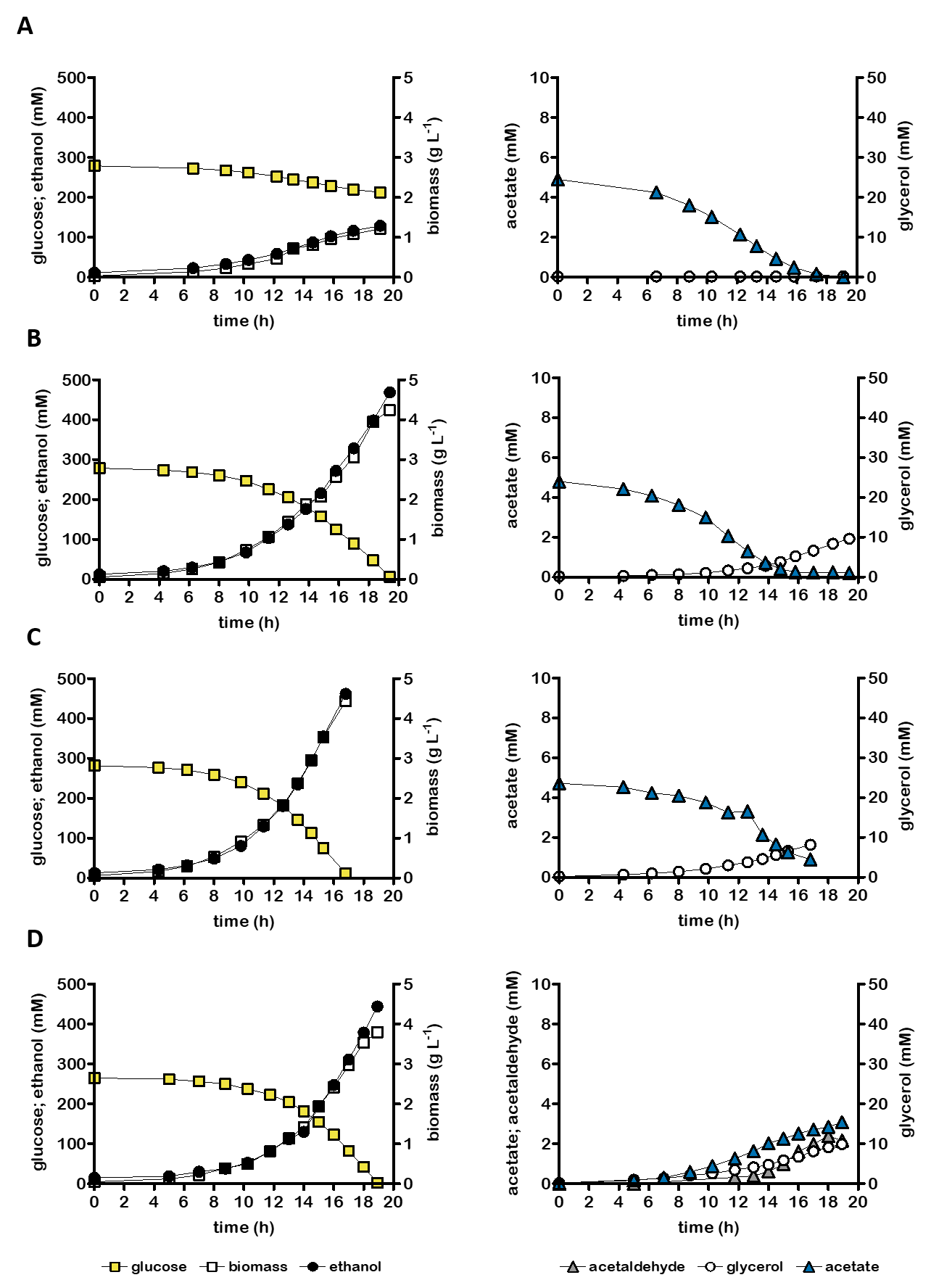

**Figure S5:** Growth, glucose consumption and product formation of anaerobic bioreactor batch cultures of S. cerevisiae strain IMS1247 (∆gpd1 ∆gpd2 ∆ald6 eutE) (**A**) and co-cultures of IMS1247 and IMX2736 (∆gpd2, non-ox PPP↑, prk, 2x cbbm, groES groEL) and IMS1247. Non-ox PPP↑ indicates integration of the overexpression cassettes RPE1, TKL1, TAL1, NQM1, RKI1 and TKL2. Cultures were grown on synthetic medium containing 50 g L^-1^ glucose (panel **D**) or 50 g L^-1^ glucose and 5 mmol L^-1^ acetate (panels **A**-**C**) and were inoculated at a ratio of ratio of 5.5±1.3 (**B**), 1.0±0.2 (**C**) or 1.3±0.4 (**D**) (inoculum ratio was estimated based on whole genome sequencing). Representative cultures of independent duplicate experiments are shown, corresponding replicate of each culture shown in Figure 5.

**Table S2: Oligonucleotide primers used in this study.** PAGE refers to polyacrylamide gel electrophoresis; DST indicates desalted.

| Primer | Sequence | purification |
| --- | --- | --- |
| 5793 | GATCATTTATCTTTCACTGCGGAG | PAGE |
| 6965 | GTGCGCATGTTTCGGCGTTCGAAACTTCTCCGCAGTGAAAGATAAATGATCGGGCAAGGACGTCGACCATAGTTTTAGAGCTAGAAATAGCAAGTTAAAATAAG | PAGE |
| 6966 | GTGCGCATGTTTCGGCGTTCGAAACTTCTCCGCAGTGAAAGATAAATGATCCCAAGAATTCCCATTATTCGGTTTTAGAGCTAGAAATAGCAAGTTAAAATAAG | PAGE |
| 6969 | GTATTTTGGTAGATTCAATTCTCTTTCCCTTTCCTTTTCCTTCGCTCCCCTTCCTTATCAAACCAATTTATCATTATACACAAGTTCTACAACTACTACTAGTAACATTACTACAGTTAT | DST |
| 6970 | ATAACTGTAGTAATGTTACTAGTAGTAGTTGTAGAACTTGTGTATAATGATAAATTGGTTTGATAAGGAAGGGGAGCGAAGGAAAAGGAAAGGGAAAGAGAATTGAATCTACCAAAATAC | DST |
